# Myosin Va Brain-Specific Mutation Alters Mouse Behavior and Disrupts Hippocampal Synapses

**DOI:** 10.1101/2020.07.08.180679

**Authors:** Swarna Pandian, Jian-Ping Zhao, Yasunobu Murata, Fernando J. Bustos, Cansu Tunca, Ramiro D. Almeida, Martha Constantine-Paton

## Abstract

Myosin Va (MyoVa) is a plus-end filamentous-actin motor protein that is highly and broadly expressed in the vertebrate body, including in the nervous system. In excitatory neurons MyoVa transports cargo toward the tip of the dendritic spine, where the post-synaptic density (PSD) is formed and maintained. MyoVa mutations in humans cause neurological dysfunction, mental retardation, hypomelanation and death in infancy or childhood. Here we characterize the Flailer (Flr) mutant mouse, which is homozygous for a *myo5a* mutation that drives high levels of mutant MyoVa (Flr protein) specifically in the CNS. Flr protein functions as a dominant-negative MyoVa, sequestering cargo and blocking its transport to the PSD. Flr mice have early seizures and mild ataxia, but mature and breed normally. Flr mice display several abnormal behaviors known to be associated with brain regions that show high expression of Flr protein. Flr mice are defective in the transport of synaptic components to the PSD and in mGluR-dependent LTD and have a reduced number of mature dendritic spines. The synaptic and behavioral abnormalities of Flr mice result in an anxiety/autism spectrum disorder (ASD)/obsessive compulsive-like phenotype similar to that of other mouse mutants with similar abnormalities. Because of the dominant-negative nature of the Flr protein, the Flr mouse offers a powerful system for the analysis of how the disruption of synaptic transport and lack of LTD can alter synaptic function, development and wiring of the brain and result in symptoms that characterize many neuropsychiatric disorders.

**SIGNIFICANCE STATEMENT:** Here we characterize a mutant mouse homozygous for a Myosin Va mutation named Flailer. The Flailer mutation generates a dominant-negative MyoVa transport motor protein that sequesters synaptic cargo and blocks synaptic transport, thereby resulting in an absence of LTD and in abnormal behaviors similar to those seen anxiety/Autism Spectrum disorders. We propose that the Flailer mutant can be used as a model to study how the absence of LTD disrupts brain connectivity and behavior. Moreover, by using the Flailer mutation together with gene editing technologies it should be possible to target specific brain areas to remove the mutation and recover MyoVa function, thereby interrogating the role of a specific brain region in the control of a particular behavior.

## INTRODUCTION

Myosin Va (MyoVa) is a plus-end filamentous-actin motor protein that is highly expressed throughout the vertebrate body, including in the nervous system (Mercer et al., 1991; Hammer and Sellers, 2011). Mutant MyoVa in humans produces an early lethal disease called Elejalde or Griscelli Syndrome Type I, which is characterized by abnormal pigmentation and severe neurological symptoms (Griscelli et al., 1978; Elejalde et al., 1979; Pastural et al., 1997; Durán-McKinster et al., 1999; Sanal et al., 2000). Similar to humans with Griscelli Syndrome, MyoVa null mice (*dilute-lethal*) die early, either *in utero* or before postnatal day (P) 21. However, rodent strains with small spontaneous *myo5a* mutations display milder phenotypes and have made possible studies of abnormal cerebellar functions (e.g. (Takagishi and Murata, 2006; Miyata et al., 2011)).

The Flailer (Flr) mouse is homozygous for a *myo5a* mutation caused by a spontaneous recombination event that produced a truncated *myoVa* gene lacking the coding region for the actin-binding “feet” of the MyoVa protein. The actin-binding domains are critical, since they allow MyoVa to “walk” toward the positive end of, and move between, actin filaments (Wagner et al., 2010; Hammer and Sellers, 2011). In the Flailer mutant, this truncated *myoVa* gene is fused to the promoter of the gene *gnb5* (guanine nucleotide-binding protein subunit beta-5) and as a consequence is expressed extensively in the CNS (Watson et al., 1994; Zhang et al., 2000; Witherow and Slepak, 2003; Lein et al., 2006; Zhang et al., 2011; *<u>Allen Brain Atlas</u>*). When the Flailer mutant protein is in a 1:1 ratio with wild-type (WT) *myo5a* protein, the Flailer protein competes with WT MyoVa for cargo by forming abnormal myosin dimers that sequester cargo but are unable to step toward the postsynaptic density (PSD) on actin, thereby diminishing MyoVa function. In the WT brain MyoVa carries many critical molecules and complexes along actin filaments in dendritic spine necks to distal spine synapses (Jones et al., 2000; Hammer and Sellers, 2011), including vesicle-bound molecules such as receptors, mRNAs translated at the synapse (Fujii et al., 2005; Yoshimura et al., 2006), and synaptic scaffolding complexes (Gerrow et al., 2006; Moutin et al., 2012; Yoshii et al., 2013). These scaffolding complexes are necessary to localize receptors and organize other molecules at the relatively isolated spine tip PSDs. Interestingly, the Flr phenotype is expressed only when the *flr* mutant gene is present in at least as many copies as the WT *myo5a* gene. More specifically, the Flr phenotype requires two copies of the *flr* mutant gene (hence appears to be recessive, although there is no second “WT” allele of the *flr* gene) in a WT *myo5a* background with two WT *myo5a* alleles, but requires only one copy (hence appears to be dominant) in a *myo5a* heterozygote with one *myo5a* null allele and one WT *myo5a* allele. These observations suggest that the *flr* mutation acts as a *myo5a* dominant-negative mutation (Jones et al., 2000).

Researchers studying MyoVa function have designed similar truncated constructs and over-expressed them in a variety of cells *in vitro* in attempts to block WT MyoVa function; however the high and scattered expression levels make the interpretation of these data difficult (Correia et al., 2008). Conversely, Flailer animals show intact cellular function and behavior that is essentially the same in every animal and thus provide an excellent model organism for the study of MyoVa function in the brain.

Previously we characterized some of the synaptic defects of the Flailer mouse visual cortex (VC), where we found LTP to be normal but no evidence of LTD (Yoshii et al., 2013). Moreover, α-Amino-3-hydroxy-5-methyl-4-isoxazolepropionic acid receptor (AMPAR) endocytosis was defective, AMPAR miniature current frequencies were significantly increased, and AMPAR to N-methyl-D-aspartate receptor (NMDAR) evoked current ratios were abnormally high in VC neurons as recorded in brain slices. Also, immunocytochemical studies of cultured VC pyramidal cells showed generally increased expression of AMPARs, which in Flr mice are located in dendritic shafts rather than in spines because of the diminished transport by MyoVa (Yoshii et al., 2013).

Here we characterize the Flailer animal using behavioral testing, biochemistry and electrophysiology. These animals show repetitive-grooming, anxiety, asocial behaviors, defective hippocampal memory, and abnormal pup vocalizations. The behaviors suggest that the Flr protein disruption of glutamate synapses produces an anxiety/autism spectrum disorder (ASD)/obsessive compulsive-like phenotype similar to those seen in mouse strains in which neuropsychiatric candidate genes have been mutated (Welch et al., 2007; McFarlane et al., 2008; Peñagarikano et al., 2011; Peca and Feng, 2012). In addition, we observed a severe deficit in the transport of synaptic components to the synapse and a lack of LTD in the hippocampus of Flailer animals. Our study highlights the important role for LTD in the etiology of mouse -- and presumably human – ASD-related syndromes and, if combined with gene editing technologies, offers a new experimental approach to identifying the brain circuits responsible for neuropsychiatric disorders.

## RESULTS

### Flailer animals show behaviors that resemble anxiety and Autism Spectrum Disorders

The *flr* gene consists of the promoter and first two exons of brain-specific *gnb5* plus part of the adjacent intron fused in-frame with an intron of *myo5a*, allowing the *gnb5* promoter to drive the expression of truncated *myo5a* and express the Flr protein (Jones et al., 2000) at levels equal to WT MyoVa in the many brain regions in which Gnb5 is expressed (Zhang et al., 2000). Gnb5 forms complexes with members of the regulators of G-protein signaling (RGS) family to control G-proteins, including mGluR6 (Rao et al., 2007). In response to light and photoreceptor glutamate release, mGluR6 activation closes a non-selective cation channel in ON-retinal bipolar cell dendrites (Morgans et al., 2007). This closure depolarizes the otherwise hyperpolarized cells.

Because abnormal expression of the Flr protein in retinal bipolar cells might seriously compromise vision, we first tested the grating acuity of four mature (P56-84) Flr and four age-matched wild-type mice with the Flr genetic background, hereafter referred to as WT_FLR_. We used scoptopic conditions, the virtual oculomotor system designed and constructed by Prusky et al. (2004), the equipment and the help of investigators in the Prusky laboratory, blind to the genotype of the mice being tested. WT animals showed average grating acuities in their right and left eyes of 0.388 and 0.389 cycles per degree (cyc/deg), respectively, while the Flr mice showed a somewhat lower acuity of 0.345 cyc/deg for both eyes, indicating that despite a slightly lower acuity the Flr mutants were sufficiently visual to detect their environments and have reasonably normal environmental stimulation of their central nervous systems.

Next, using adult (P56-84) male Flr and WT mice, we characterized the behaviors displayed by the animals. In all cases both training and testing took place within the first 3 hours of the dark cycle. To determine the anxiety levels shown by Flailer animals, we performed three tests. The most salient feature of Flr behavior as compared to that of the WT_FLR_ strain was the frequency and completeness of their spontaneous bouts of self-grooming (Figure 1A; Movie 1&2). Individual bouts could last up to 6 minutes and the sequence of this grooming behavior in Flr was nearly identical with the normal syntactical behavior described by Berridge and Whishaw (1992) for rats. This behavior is regularly associated with high anxiety and autism spectrum disorders (ASDs) in mice. Because this behavior involved full-body grooming by Flr, it did not cause the loss of head-fur described for several mice strains carrying SAPAP or Shank3 deletions (Welch et al., 2007; Peça et al., 2012). In addition, we made use of the elevated plus-maze (McNaughton et al., 2008) and found no significant difference in the entries into the open arms between Flr and WT_FLR_ mice (Figure 1B). However, Flr mice spent significantly less time in the open arms compared to WT mice, suggesting increased anxiety (Figure 1C). Rearing in the closed arms, considered an exploratory behavior, was also significantly decreased in Flr versus WT_FLR_ (Figure 1D), but frequencies of head dips in the open arm, considered a risk-taking behavior (McFarlane et al., 2008; McNaughton et al., 2008), were similar in Flr and WT_FLR_ (data not shown). Lastly, we tested the animals in a light/dark paradigm (Costall et al., 1989). We found that Flr animals spent less time in the bright field (Figure 1E) and showed a higher latency to enter the dark field (Figure 1F) than WT_FLR_ animals. Together our results demonstrate that Flailer animals show strong anxiety and repetitive behaviors similar to those of other ASD and anxiety mouse models.

**Figure 1.**
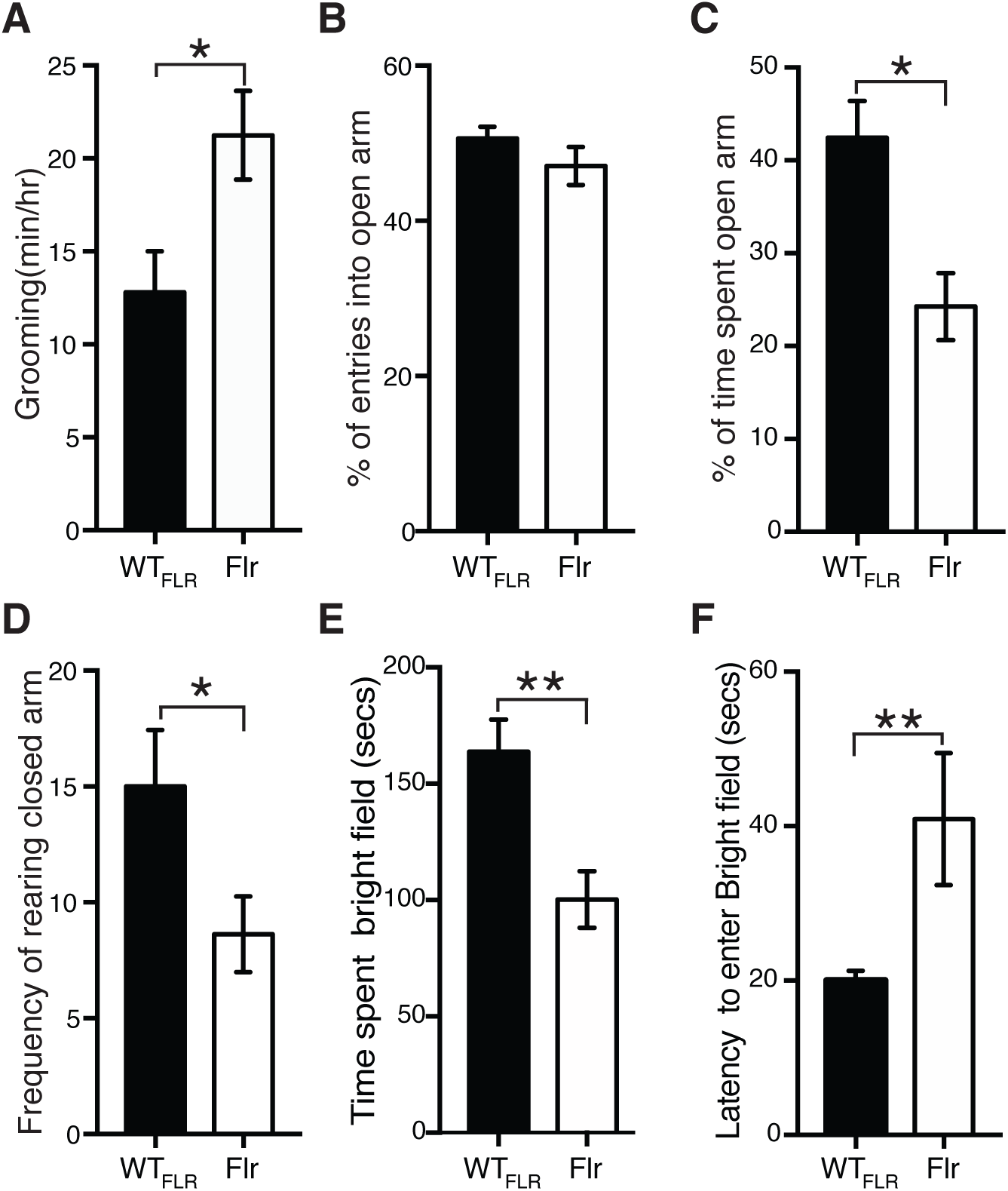
Flailer mice show anxiety-like behaviors. (A) Quantification of the time spent grooming (n = 6 for both Flr & WT_FLR_ ; Movie 1 & 2). (B-D) Elevated plus maze test to test entries into the open arms (B), time spent in the open arms compared (C), and rearing (D). (E) Light and Dark test to measure time spent in bright field (E) and latency to enter bright field (F) (n = 11 for both WT_FLR_ & Flr). Unpaired t-test was used for all the statistical analysis unless specified. *p ≤ 0.05, **p ≤ 0.001, ***p ≤ 0.0001. Error bars represent ±SEM.

Another characteristic behavior of ASD phenotypes is the lack of aptitude to perform social interactions. To determine Flailer’s ability to interact socially we used the three-chamber social interaction test (Bolivar et al., 2007; McFarlane et al., 2008; Defensor et al., 2010). In the three-chamber social interaction test WT_FLR_ and Flr mice spent approximately the same amount of time with the novel mouse (NM) and less but insignificantly different times in the chamber with the novel object (NO) (Figure 2A). Interestingly, when we reviewed the behavior of the mice in the chamber with the NM in detail and blinded to strain difference, we found that Flr spent significantly less time than the WT adjacent to the NM (Figure 2B) and if adjacent to the NM, Flr spent most of its time grooming (Movie 3&4). To gain insight into this behavior, we tested for direct interactions by quantifying social sniffing between animals. Flr/Flr or WT_FLR_/WT_FLR_ pairs were contained in a clear rectangular chamber (7 cm × 14 cm ×30 cm) that required some physical contact between the animals (Defensor et al., 2010). Flr pairs showed a significant reduction in nose-to-nose (NN) sniffing compared to WT pairs. There were also significant increases among Flr pairs compared to WT pairs in nose-to-anogenital (NA) sniffing and in nose-to-head (NH) sniffing. The latter might reflect attempts of Flr to avoid accidental nose-to-nose contact, a possibility consistent with the additional finding that crawl-over (CO) behavior by the recipient mouse, which avoids NN and NH sniffing, was also significantly increased in Flr pairs compared to WT pairs (Figure 2C). To establish that general olfactory behavior was not disrupted in Flr, animals were tested in isolation for non-social sniffing behavior. Both WT_FLR_ and Flr spent similar amounts of time sniffing a water saturated wick, and, in a separate exposure, a vanilla extract saturated wick, suggesting that general olfactory behavior of the strains was not perturbed (data not shown). Our data suggest that Flr/Flr pairs actively avoid head region close contact (Figure 2C). To test this hypothesis, we placed Flr/WT_FLR_ pairs together in the same close proximity box, reasoning that if WT mice_FLR_ persisted in head region contact Flr might demonstrate a more obvious avoidance behavior. We found that the Flr member of each pair avoided the WT_FLR_ sniffing near the head by turning away abruptly, or actively, with forepaws pushing the WT_FLR_ away (Figure 2D; Movie 5&6). Moreover, Flr animals also engaged in significantly more instances of grooming-onset during this 2 min interval compared to WT_FLR_ (Figure 2D) suggesting an increase in anxiety. Together our data indicate that Flailer animals display severe social deficits, avoiding repeatedly and strongly contact with other animals.

**Figure 2.**
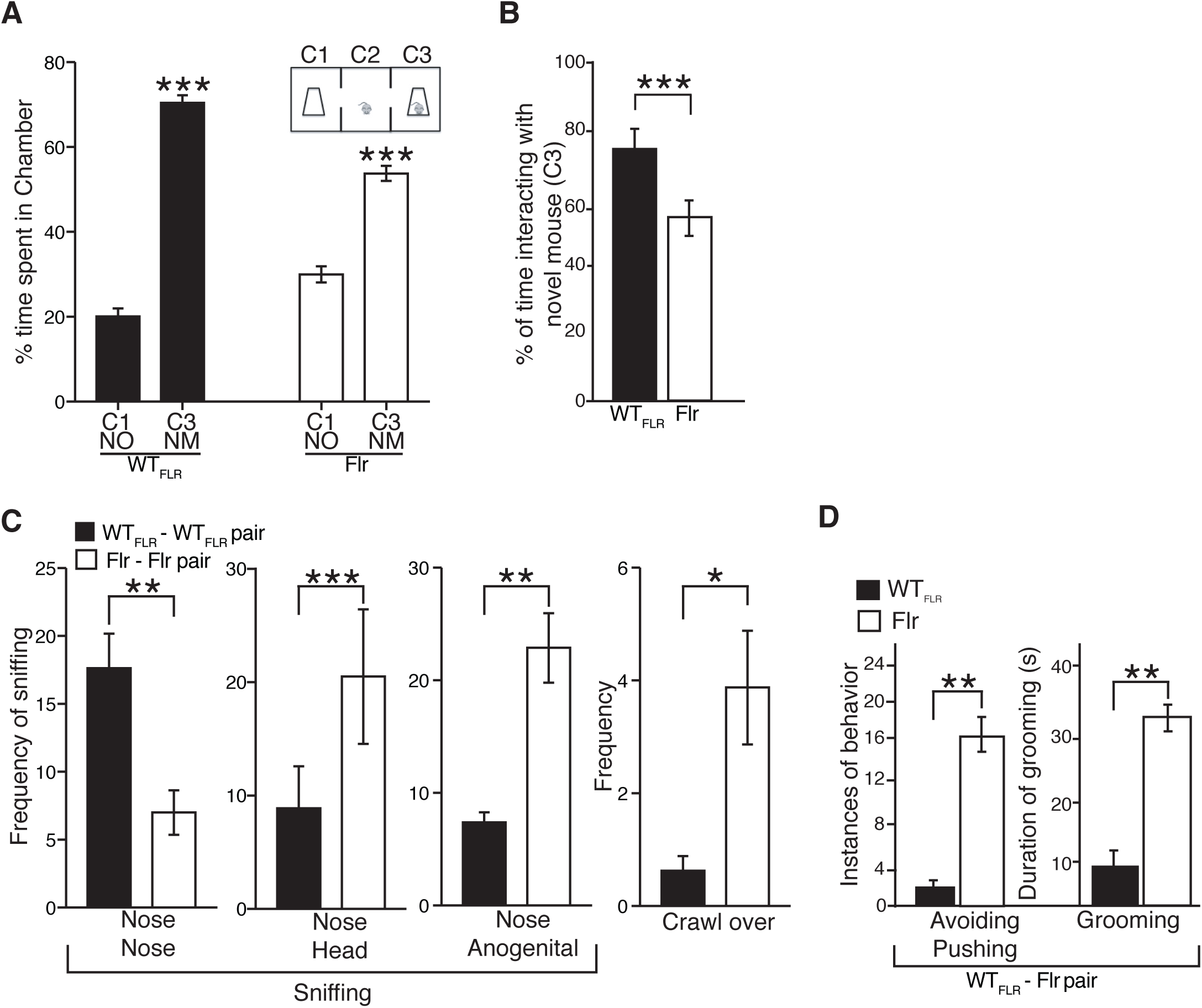
Atypical social interactions in Flr mice. (A) Three-chamber social approach test where a caged novel mouse (NM) in chamber 3 (C3) and a novel object (NO) in chamber 1 (C1) were placed (n = 17 for both WT_FLR_ & Flr). (B) Percentage of time spent interacting with NM (sniffing and close proximity to the NM cage) (Movie 3 & 4). (C) Social proximity test with Flr-Flr and WT_FLR_-WTFLR pairs. The behaviors measured were nose tip-to-nose tip sniffing (with/without contact) (NN), nose-to-head sniffing (NH), nose-to-anogenital sniffing (NA) (n = 20 for both WT_FLR_ & Flr). (D) Social proximity test with Flr-WT_FLR_ pairs Flr to quantify avoiding/pushing and grooming behaviors (Movie 5 & 6).

To test if memory formation in Flailer animals is disrupted, we conducted the Morris water maze test and fear conditioning tests. For spatial memory each Flr or WT_FLR_ mouse was trained during repeated 90 sec periods to find a submerged platform in an environment with distinct visual cues until all mice found the submerged platform with essentially the same efficiency as indicated by shortened swim distances compared to their first exposure (Figure 3A Trials 1 and 2). Retention of the platform’s position was tested 24 hrs later. In this test Flr mice failed to retain the memory of the platform position: they spent significantly less time in the quadrant containing the platform and covered significantly longer swim distances to reach the platform compared to WT_FLR_ (Figure 3A, Test). In addition, we tested memory formation dependent on either hippocampus or amygdala using the fear conditioning paradigm based on context or cue, respectively. Each individual mouse was placed in a test box with a floor that could deliver a short electric shock (MED Associates, St. Albans, VT). All mice explored the environment with little or no freezing. For hippocampal-dependent memory formation, in a second individual introduction to the box, every mouse received a 2 sec, 0.6 mA electric shock to its feet (Figure 3B Context Test, Baseline). When each mouse was subsequently re-introduced to the box, Flr mice showed a significant decrease in freezing compared to WT_FLR_ (Figure 5C; Context Test), suggesting defects in hippocampal dependent learning/memory. To test for amygdala-dependent memory formation caused by fear, the apparatus consisted of the same box plus an 85 db burst of noise delivered before the shock. Both Flr and WT_FLR_ mice were first tested with a noise stimulus and a shock. Both strains of mice responded with freezing, and no significant difference was observed between Flr and WT_FLR_ (Figure 3B Cued Test, Baseline). Animals were reintroduced to the box after approximately 2 hr and given the sound cue without the shock. Both Flr and WT_FLR_ spent approximately the same time freezing, indicating that cued-learning was not significantly different between WT_FLR_ and Flr (Figure 3B; Cued Test). Together, our data indicate that Flr has severe hippocampal-dependent memory formation deficits and that fear-conditioned memory formation dependent on the amygdala is not affected.

**Figure 3.**
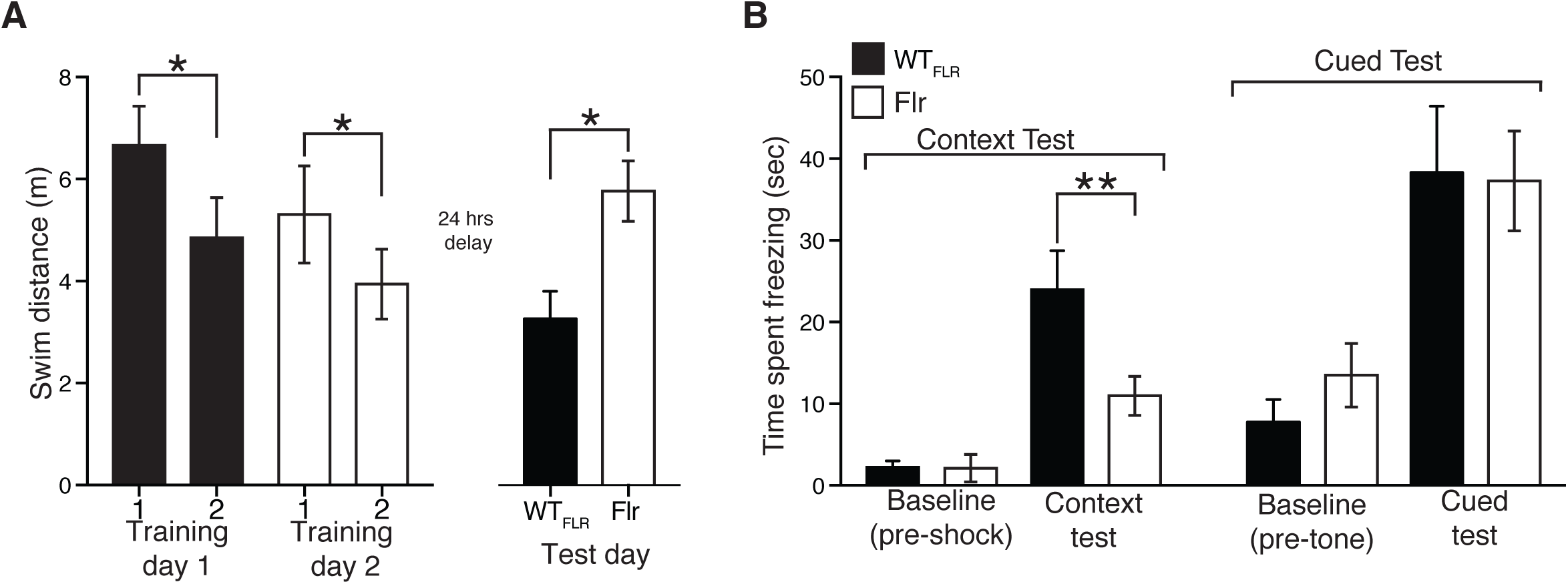
Flr mice show deficient hippocampus dependent-spatial memory formation. (A) Morris water maze on WT_FLR_ and Flr animals on training and test days (n = 9, 8 trials per animal on each training day). (B) Context-dependent and cued fear (85 dB noise) conditioning paradigms of WT_FLR_ and Flr animals. Paired T test was used for statistical analysis.

**Figure 4.**
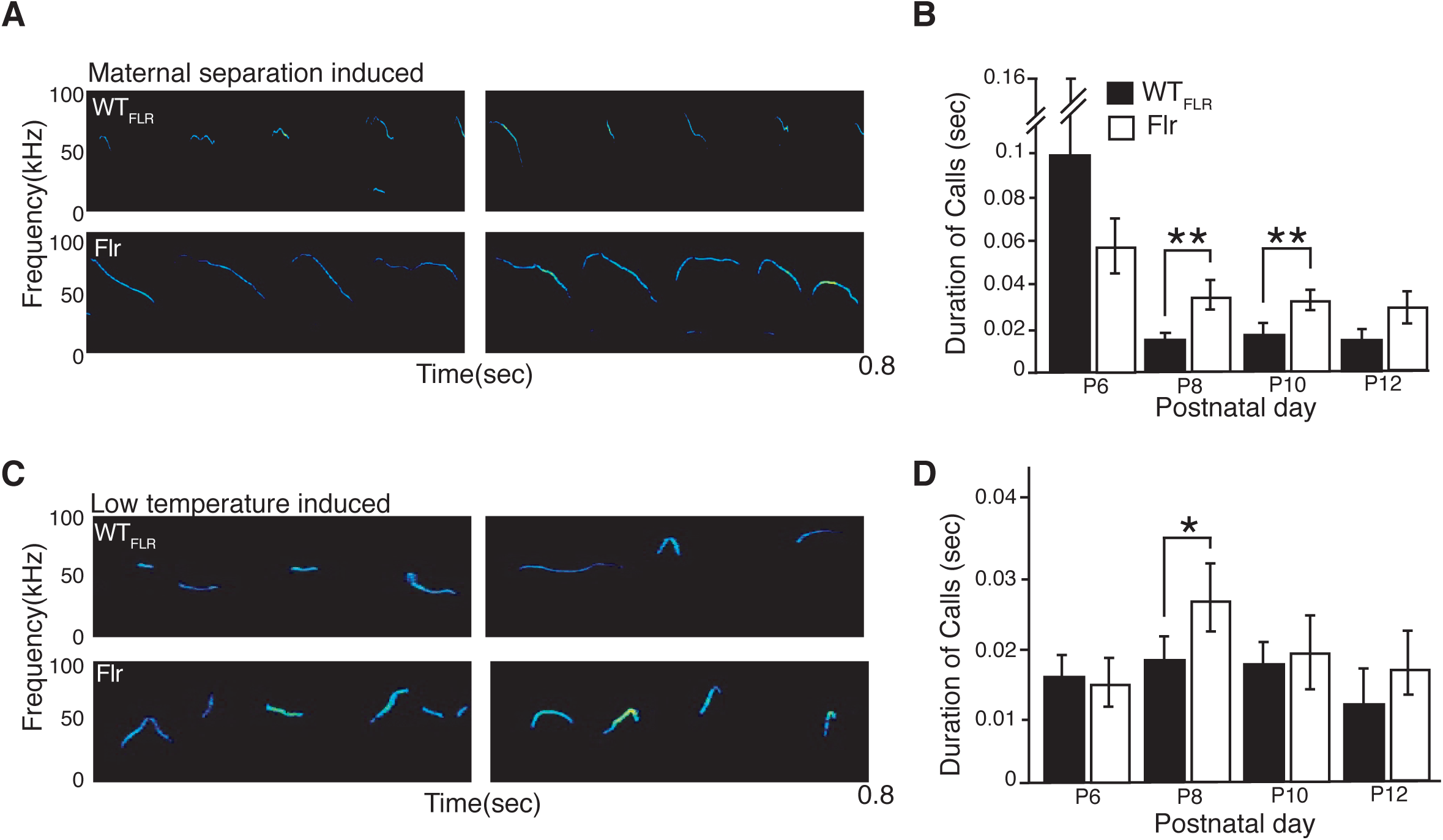
Communication deficits in Flr mice. (A) Representative spectrograms of a P8 pup during maternal separation. (B) Maternal isolation induced ultrasonic vocalization (USV) on P8 and P10 (n =10 for both WT_FLR_ & Flr). Flr showed statistically significant increase in total duration of calls. (C) Representative spectrogram of P8 pup USV during cold-temperature induced vocalization (D) Cold temperature induced USV (n = 10 for both WT_FLR_ & Flr). Flr showed statistically significant increase in total duration of calls on P8.

**Figure 5.**
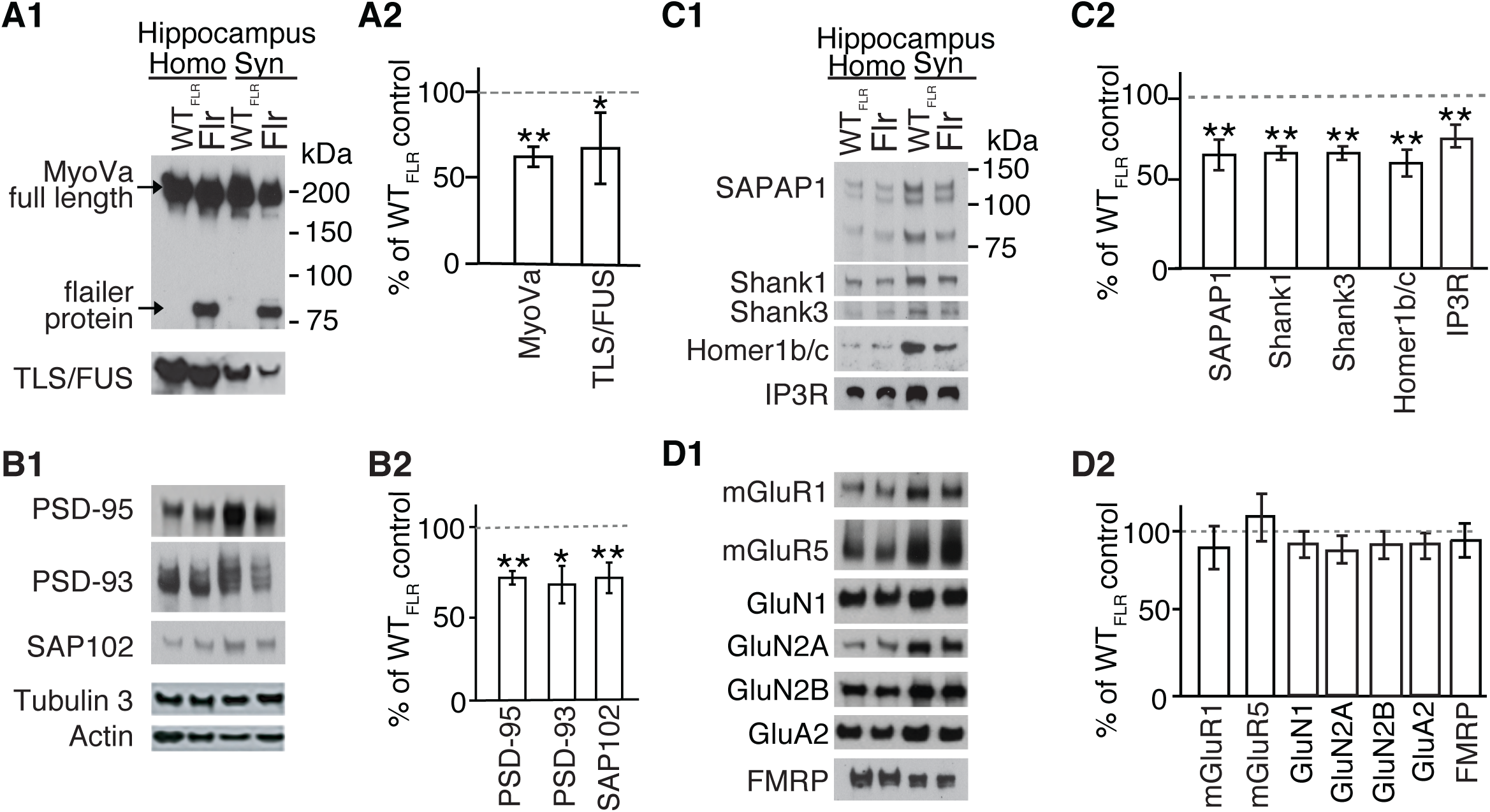
Synaptic transport of proteins to PSD is altered in Flr mice. WT_FLR_ and Flr hippocampal synaptosomal extracts were used for quantitative western blot analyses of synaptic proteins. (A1&A2) Levels of full-length MyoVa and TLS/FUS were reduced in Flr extracts. (B1 & B2) Expression levels of major MAGUKs (PSD-95, PSD-93, SAP102) and (C1 & C2) SAPAP1, SHANK1, SHANK3, Homer1b/c and IP3R were significantly reduced in Flr. (D1 & D2) mGluRs, NMDAR, AMPAR subunit levels and FMRP were not changed in Flr when compared to WT_FLR_. Homogenate (Homo) and synaptosome (Syn) was prepared from whole hippocampus. Protein levels in Flr were quantified and plotted as % of WT_FLR_. n = 3 samples per group/ each sample pool 3-5 animals.

Another characteristic behavior observed for ASD mouse models is an increase in the duration of ultrasonic vocalizations (USVs) (Wöhr et al., 2011). This abnormal vocalization has been observed in BTBR T+tf/J and *Shank1* mutants, two models of ASD (Scattoni et al., 2008; Wöhr et al., 2011). To test if Flr mice also display this ASD-linked phenotype, we evaluated changes in USVs in Flr and WT_FLR_ pups after separating individuals from their mothers and littermates or after exposing isolated pups to low-temperatures. Under separation conditions, all mouse pups produced infrequent and relatively long calls starting on P6. However, when the same pups were tested again at P8, and P10, isolated from their mothers and littermates, the duration of the Flr pup calls were invariably longer than those of WT_FLR_ (Figure 4 A, B). In addition, calls induced by low temperature isolation also had longer durations in Flr than in WT_FLR_ but only at P8 (Figure 4 C, D). Taken together, out data show severe behavioral deficits of Flr animals, which correlates with an anxiety-ASD-like phenotype.

### Flailer animals are deficient in the transport of synaptic components to the post-synaptic density

Next, we asked whether the severe changes in animal behavior observed in Flr are caused by the diminished transport of synaptic components to the PSD. The imaging analyses of Gerrow et al. (2006) and Moutin et al. (2012) demonstrated that glutamate synapse scaffolding molecules -- SAPAP (GKAP), Shank and the membrane-associated guanylate kinase (MAGUK) PSD-95 -- travel as a complex up dendritic shafts and then up spine necks to the PSDs at the spine tip. In Flr, a significant amount of synaptic cargo gets transferred to a dimer of the truncated MyoVA cargo-binding domain (e.g. the Flr protein). These abnormal myosins, lacking the end-feet with ATP hydrolyzing heads of WT MyoVA, cannot step along the actin filaments to the PSD. However, it is possible than more than one MyoVa motor attaches to a single transport vesicle and moves in a straight line along an actin filament (Hammer and Sellers, 2011). This would account for the presence of Flr protein (∼85 kD) in Flr mouse synaptosomes (Figure5 A1). However, WT MyoVA carrying some of the scaffolds attached to the Flr protein cargo could be slowed down or unable to move to the PSD. This would account for the decrease in WT MyoVA found in Flr synaptosomes (Figure 5 A1, A2).

To characterize changes in transport of synaptic components in Flr neurons, we compared the levels of proteins between WT_FLR_ and Flr synaptosomes using quantitative western blotting for 15 synaptic proteins: 1. Known MyoVA synapse-associated cargos: the mRNA binding protein TLS/FUS (Yoshimura et al., 2006), and the IP3R (Takagishi et al., 1996; Miyata et al., 2011), and Fragile X Mental Retardation Protein (FMRP). 2. The major mature glutamate synapse scaffold complex, namely PSD-95, SAPAP and Shank (Gerrow et al., 2006; Moutin et al., 2012) and molecules closely related to PSD-95, namely the MAGUKS PSD-93 and SAP102 (Nithianantharajah and Grant, 2013) and Homer1c known to be directly associated with Shank at synapses (Tu et al., 1999). 3. The major glutamate receptors or glutamate receptor subunits, namely mGluR1, mGluR5, GluN1, GluN2A, GluN2B and GluA2. Nine of these proteins TLS/FUS (Figure 1 A1, A2), PSD-95, PSD-93, SAP102 (Figure 5 B1, B2), SAPAP, Shank1, Shank3, Homer 1c, IP3R (Figure 5 C1, C2) showed significant reductions in their expression in Flr synaptosomes compared to synaptosomes from WT mice. However, all glutamate receptors and receptor subunits tested (mGluR1, mGluR5, GluN 1, 2A, 2B, A2) as well as the FMRP protein, were not significantly different in synaptosome levels between WT_FLR_ and Flr (Figure 5 D1, D2). Together, we found a significant reduction on the expression of the scaffolding complexes in synaptosomes of Flr neurons. Conversely, all receptors subunits studied did not show changes in the expression of proteins in Flr synaptosomes. This could be caused by the abnormal clustering of receptors at dendritic shafts (Yoshii et al., 2013) that might have rounded as vesicles and precipitated along with synaptosomes. The changes observed in synaptic composition because of the deficient transport by MyoVa could in fact explain the behavioral deficits observed in Flr mice shown above.

### mGLuR-dependent Long-Term Depression (LTD), but not Long-Term Potentiation (LTP), is impaired in Flailer mutants

We have previously shown that >6 month old Flr animals have impaired LTD in visual cortex slices, indeed showing a small LTP response in its place (Yoshii et al., 2013). Many studies have proposed that synaptic plasticity phenomena such as LTP and LTD are the main mechanisms that control memory formation (Nabavi et al., 2014). Here we have shown that Flailer animals have impaired hippocampal-dependent memory formation (Figure 3), which could be a consequence of the lack of synaptic plasticity phenomena such as LTD. For this reason, we asked if LTP and LTD at the Shaffer Collateral (SC) to CA1 pyramidal synapse (SC-CA1) is altered in Flr mice.

To study fLTP in CA1 pyramids, we induced LTP by SC stimulation using three different protocols: Two 1 sec 100 Hz bursts at 20-sec inter-burst intervals, involving both NMDARs and some L-type Ca^2+^ channels; a single 1 sec 25 Hz burst known to be dependent only on NMDARs (Cavus and Teyler, 1996); and saturation LTP, with six 100 Hz stimuli at 5 min intervals. In all three protocols used to induce LTP, responses between Flr and WT_FLR_ animals were indistinguishable, showing no defects in Flr animals (Figure 6 A-C, n= number of slices/ number of mice). Conversely, two forms of LTD occur at SC-CA1 synapses. The first group requires mGluR5 activation of Phospholipase C to generate IP3 and diacylglycerol (DAG). IP3 activates the IP3R on SER to release Ca^2+^ into spine cytoplasm and DAG triggers local protein synthesis (Huber et al., 2000). In WT rodents this form of LTD is induced by stimulating the SC-CA1 inputs with paired-pulses separated by 50ms and given at 1 Hz in the presence of the NMDAR antagonist AP5 (Kemp and Bashir, 1999) or by application of the metabotropic Group1 mGluR agonist DHPG (Palmer et al., 1997). Both stimulation protocols produced normal mGluR5-dependent LTD in WT_FLR_ mice (Figure 6 D, E). In contrast, Flr mice showed a small LTP in response to the paired-pulse stimulation protocol and no LTD in response to the agonist DHPG (Figure 6 D, E). To determine the dependency on protein synthesis, we used the protein synthesis inhibitor anisomycin. In WT_FLR_ (control) mice mGluR5 LTD was blocked, showing a response similar the induction of LTP (n=6 slices from 5 mice; data not shown). Conversely, Flr animals showed no mGluR5 LTD and a small LTP regardless of whether anisomycin was present (n=15 slices from 13 mice without anisomycin; n= 5 slices from 5 mice with anisomycin; data not shown).

**Figure 6.**
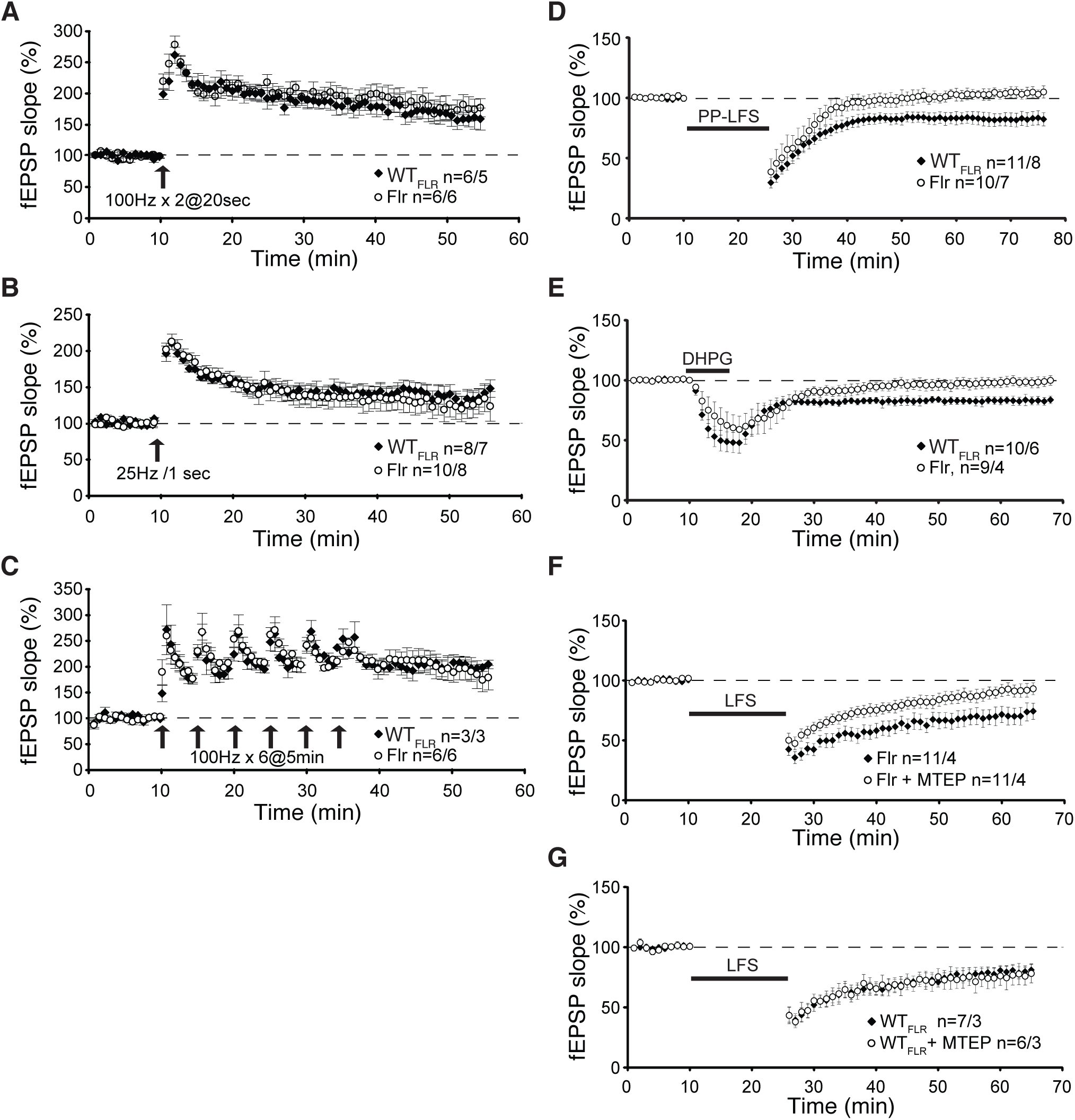
Absence of mGluR-dependent LTD at hippocampal SC to CA1 synapses in Flr mice. Field potentials (fEPSPs) were recorded in the stratum radiatum of CA1 by delivering stimulation to Schaffer collateral pathway (SC) in both WT_FLR_ and Flr mice. LTP was induced by (A) Two 1 sec 100 Hz bursts at 20 sec inter-burst interval of 1s duration and 20 s inter-train interval (arrow); (B) A single 1 sec 25 Hz burst (arrow); (C) Saturating induction, six 1 sec 100 Hz bursts at 5 min inter-burst intervals (arrows). (D) mGluR-dependent LTD was induced In the presence of 50 µM D-AP5, PP-LFS (Kemp and Bashir, 1999) (black bar). (E) Chemical induction of LTD using the mGluR agonist DHPG (50 µM) applied for 5 min (black bar). (F) Low frequency stimulation (LFS) LTD (900 stimuli at 1 Hz, black bar) induced NMDAR-dependent LTD in Flr that is inhibited by MTEP (4 µM). (G) LFS of WT_FLR_ mice was not affected by the specific mGluR antagonist MTEP.

In contrast to mGluR5 LTD, NMDAR-dependent LTD induced by 900 stimuli at 1 Hz in Flr animals appeared normal (Figure 6F black squares), a finding also reported by Schnell and Nicoll (2001) based on recordings from young dilute-lethal mice. However, our data support the hypothesis that since some WT MyoVA could reach Flr synapses, there might be low, but sufficient, Ca^2+^ release through residual IP3R and mGluR5 expression/activity (Figure 5) at the synapses of some cells to facilitate NMDAR LTD. Consequently, we tested for an effect of residual mGluR5 facilitation of NMDAR-LTD by applying the highly specific mGluR5 antagonist MTEP (Cosford et al., 2003) to block any remaining mGluR5 activity in slices from Flr mice. We found that MTEP significantly reduced NMDAR-dependent LTD in these Flr brain slices (Figure 6F, open circles). This effect was not observed in WT_FLR_ mice of the same age (Figure 6G, overlapped data). This observation precludes any involvement in NMDAR-LTD of residual Ca^2+^ release from SER or tyrosine phosphorylation of NMDARs by Group1 mGluRs (Heidinger et al., 2002). Instead, these results indicate a mitigating effect of mGluR5 on the disruption of NMDAR-dependent LTD in Flr, resulting from the loss of normal scaffolding proteins.

To confirm our conclusion that the loss of LTD is dependent on mGluR activity, we used a positive allosteric modulator (PAM) of mGluRs, (3-cyano-N-(1,3-diphenyl-1H-pyrazol-5-yl) benzamide) or “CDPPB” (Kinney et al., 2004). In Tuberous Sclerosis Tsc2^+/-^ mutant mice, a well-known mental retardation, epilepsy and ASD animal model, the application CDPPB is able to restore normal mGluR-LTD (Auerbach et al., 2011). Consequently, we tested whether incubation of Flr hippocampal slices with CDPPB could rescue mGluR5-LTD defects. We found that pretreatment with CDPPB either applied to the slices for 35 min prior to recording (Figure 7B) or applied for 30 min before and then during recording (Figure 7C) did not restore defective mGluR-LTD in Flr (Figure 7 A-C). Together our data show that in hippocampal slices Flailer animals are able to produce normal LTP but that mGluR5-dependent LTD is severely impaired.

**Figure 7.**
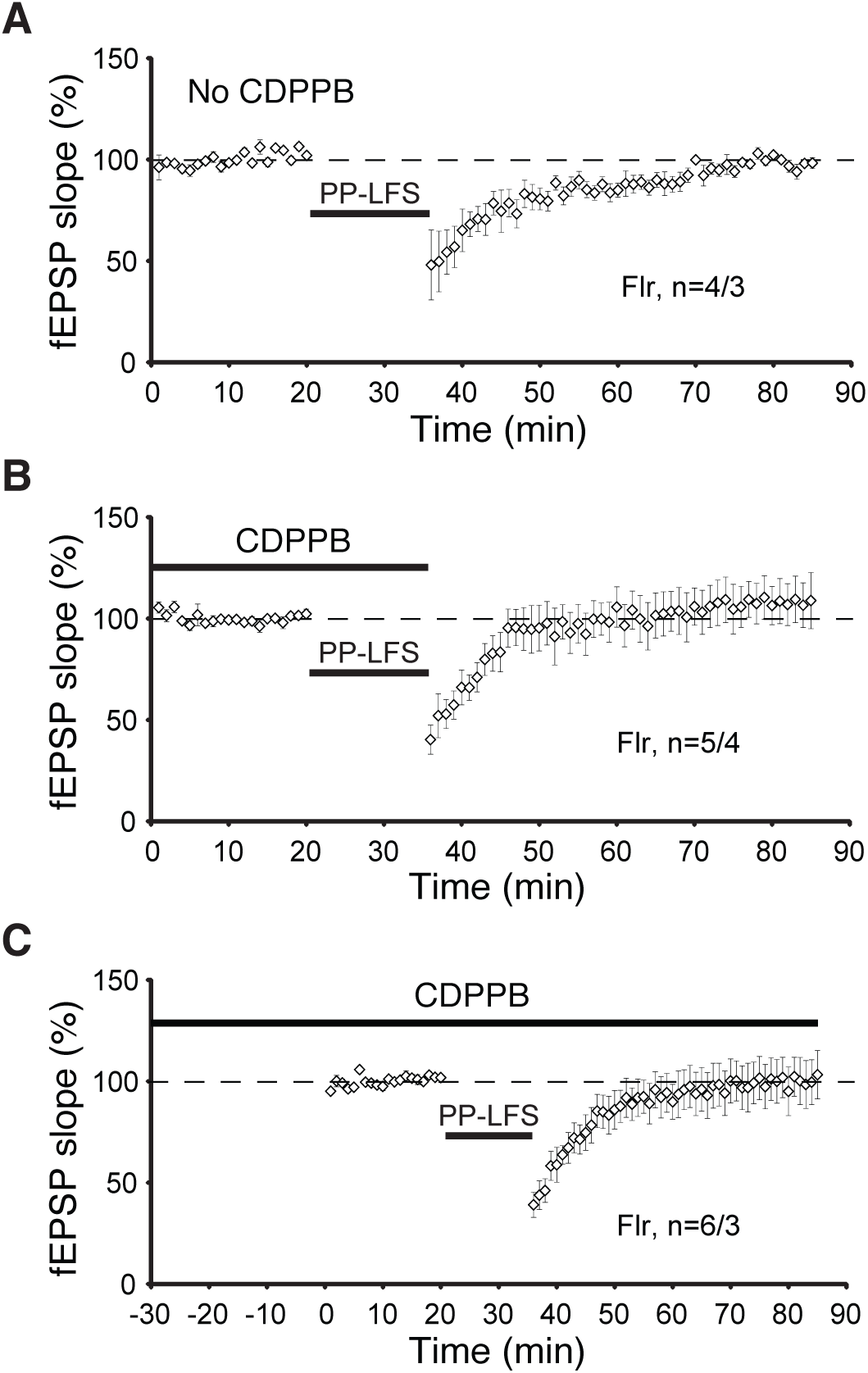
mGluR-dependent LTD is not rescued by mGluR5 agonist CDPPB in Flailer mice. (A) LFS does not induce mGluR-dependent LTD in Flailer animals. (B) CDPPB (10 µM) was added to the bath 30 min before and during LFS induction protocol for LTD. (C) CDPPB was added to the bath throughout the experiment, 30 min before, during and after LFS mGluR-dependent LTD protocol. All recordings above were performed in the presence of 50 µM D-AP5.

### Flailer mutants have a reduced number of mature dendritic protrusions in CA1 pyramidal neurons

In many human and mouse neurodevelopmental disorders, it has been shown an abnormal increase in spine density and filipodia shaped spines (Nimchinsky et al., 2001; Hutsler and Zhang, 2010; Penzes et al., 2011; Kulkarni and Firestein, 2012; Peça et al., 2012; Yoshii et al., 2013). To determine changes in spine morphology and density, we examined CA1 pyramidal cells, facilitated in Flr because the Flr strain used for this analysis was backcrossed to homozygosity after mating Flr with Thy1-GFP mice (Feng et al., 2000). Five apical dendritic segments, ∼15-20 µm in length and starting ∼150 µm from the neuron’s soma, from each of 8 mature Flr males and 8 mature WT_FLR_ males, were chosen for quantitative analyses. All scoring of protrusion type was done blinded to the phenotype of the mouse. Flr dendritic shafts had many more irregular, frequently thin protrusions than WT_FLR_ neurons (Figure 8A). We found a significant increase in Flr as compared to WT_FLR_ in filopodia density (Figure 8B, 8A arrows; p = 0.02); no difference in thin spines (Figure 8B, 8A asterisk) and more mushroom spines in WT_FLR_ compared to Flr (Figure 8B, 8A arrow heads; p = 0.03). Together our data suggest that defective transport of synaptic proteins leads to deficits in spine density and maturation in Flr mice.

**Figure 8.**
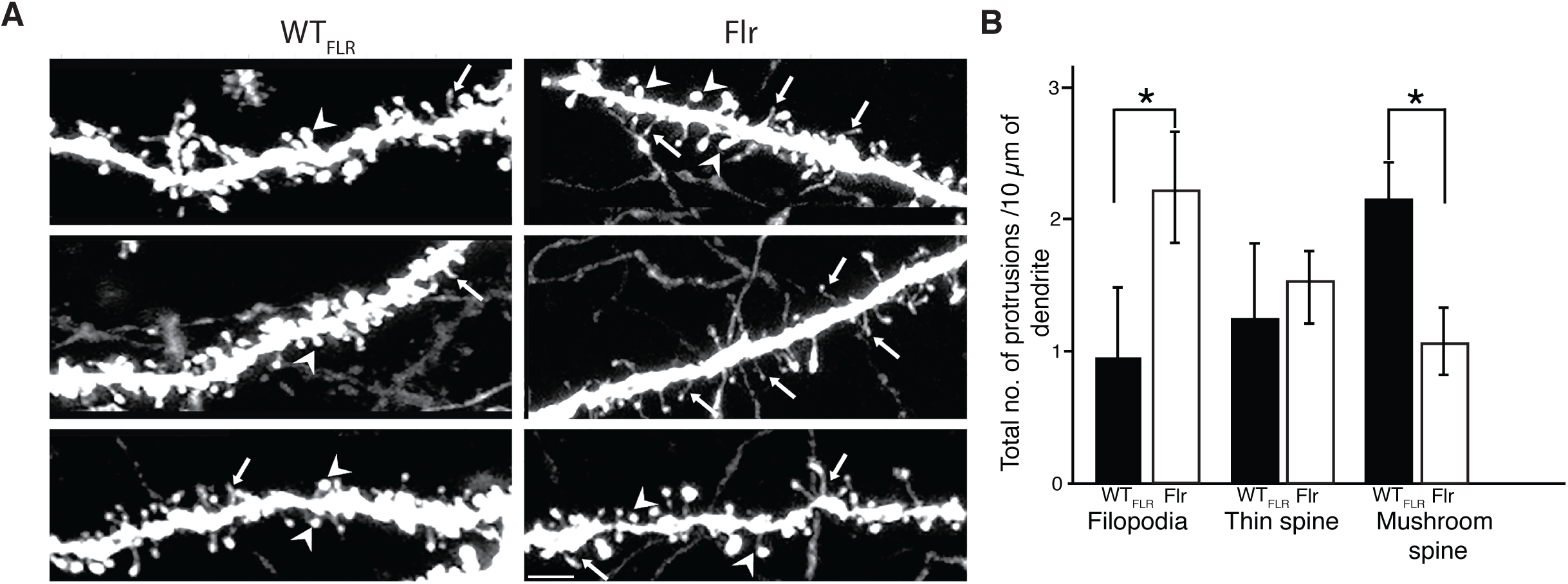
Flr shows increased number of filopodia and decrease in mushroom spines in apical dendrites of hippocampal pyramids. (A) Representative confocal images of apical dendritic shafts showing filopodia (arrows), thin spines (asterisk) and mushroom spines (arrowheads) in both Flr & WT_FLR_. (B) Quantification of the number of filopodia and mushroom spines in Flr and WT_FLR_ (n = 8 animals/genotype and 40 segments of length 15-20 µm). The spine analysis and reconstruction was done using autoneuron of neurolucida and contrast increase was done using imageJ and adobe CC Scale bar = 10 µm

## DISCUSSION

Myosin Va has many functions in the body, including within the nervous system. Myosin Va is a prominent and versatile motor for delivering cargo close to its target zone (Langford, 2002; Yoshimura et al., 2006). Our results report the consequences of disrupting this motor’s critical role in establishing proper pre-to post-synaptic transfer of neurotransmitter-mediated nervous system information. In all neurons with post-synaptic densities (PSDs) on dendritic spine tips MyoVa delivers ribonuclear particles for local synthesis of post-synaptic proteins (Yoshimura et al., 2006) and a variety of cargos associated with spine PSD components, such as smooth ER, TrkB, mRNA/protein complexes, the molecule PSD-95 and additional scaffolding. All are crucial to normal glutamate depolarization at spine PSDs and post-synaptic cell activation (Naisbitt et al., 2000; Yoshimura et al., 2006; Wagner et al., 2010; Sui et al., 2015). Consequently, in Flr mice homozygous for a dominant-negative MyoVa mutation, the disruption of the transport of these synaptic components causes severe deficits in synaptic function and behavior.

Many analyses of glutamate PSD structure have established the scaffolding system at glutamate synapses (Sheng and Hoogenraad, 2007). Electrophysiological abnormalities such as disruptions in mGluR LTD and changes in dendritic protrusions in the Flr hippocampus as well the abnormal behaviors of Flr mice indicate that this animal’s truncated MyoVA causes a phenotype similar in some respects to those of mice that have been genetically engineered to have one of their native scaffolding genes altered to mimic mutations found in humans with ASD or afflicted with other neuropsychiatric syndromes (Welch et al., 2007; McFarlane et al., 2008; Peñagarikano et al., 2011; Peça et al., 2012; Peca and Feng, 2012; Ronesi et al., 2012; Sato et al., 2012). These genes include those encoding proteins that are known cargos of the MyoVA transport system, such as SAPAP, Shank, and PSD-95. One example of a shared abnormal behavior is the self-destructive repetitive behavior that causes mice to lose fur or bloody themselves with compulsive grooming of one body region. This behavior is seen with SAPAP and some SHANK mutations and can be eliminated if SAPAP3 is replaced in the striatum (Welch et al., 2007). Mouse models with Shank mutations produce autism-syndrome-like behaviors (Wang et al., 2011; Wöhr et al., 2011; Peça et al., 2012; Sato et al., 2012; Jiang and Ehlers, 2013). Mutations in PSD-95 itself have been seen in some people with ASD (Feyder et al., 2010; Tsai et al., 2012; Cao et al., 2013). In addition, proteins bound at the synapse by these major scaffolds at the PSD, such as multimerized or long Homers 1b or 1c (Ronesi et al., 2012) and neuroligins (Yu et al., 2011; Baudouin et al., 2012; Wöhr and Scattoni, 2013), are also implicated in human autism or autism-like behaviors in mice. The Flr mouse is significant because it is the only *myo5a* rodent mutant in which the *myo5a* mutation is brain-specific and the only *myo5a* mutant that survives throughout adulthood, hence able to show abnormal behavior. In addition, the expression pattern of the *flr* mutation’s promoter (*gnb5)* has been mapped in the mouse brain (Allen Mouse Brain Atlas), facilitating identification of the contribution of mutation-specific brain regions and to particular behaviors.

The issue of whether the Flr mouse specifically models humans with ASD is complex and impossible to answer at present, since many of the interactions that cause ASD and other neuropsychiatric diseases are unknown. Many mouse mutants with the single mutations seen in autism families show the anxiety, avoidance and disruption of normal vocalizations seen in Flr mice (Figure 1-2-4) (Welch et al., 2007; McFarlane et al., 2008; Peñagarikano et al., 2011; Peça et al., 2012). In humans subsequently diagnosed as being on the autism spectrum, early onset seizure with subsequent recovery is often present (Berg et al., 2011; Jeste, 2011), and autistic children are frequently asocial, anxious, show repetitive behaviors and become socially awkward as adults. The Flr mouse shows similar characteristics, but its repetitive behavior is elaborate, frequently including a full-body grooming sequence. Such behavior has not been seen in mouse models carrying single human ASD-associated mutations and might reflect an unknown whole-body somatosensory defect. However, we suggest that such a whole-body sensorial defect is unlikely, since Flr/Flr pairs are able to engage in social interactions with their littermates showing excessive nose-to-anal sniffing and contact (Figure 2C). In addition, the behavior of a Flr mouse in a confined area with a WT mouse is aggressive, as if touching in the head area were sufficiently threatening to a Flr mouse that it needs to push the WT mouse away with its forepaws when approach is near its head (Movie 5&6). This view is consistent with our findings showing severe anxiety behaviors displayed by Flr mice in the plus maze, and also with the repetitive grooming that occurs whenever Flr mice are in an unfamiliar environment (Figure 1).

The high expression of *gnb5* in dorsal hippocampus motivated us to test Flr and WT_FLR_ spatial memory using the water maze and the assay for context-dependent freezing. Flr mice failed in both of these tests that are dependent on spatial memory mediated by the in dorsal (CA1) hippocampus (Morris, 1984; Tsien et al., 1996). Conversely, our experiments using a cued fear-conditioning paradigm dependent mainly on the amygdala (McHugh et al., 2004) showed no difference between Flr and WT_FLR_ animals, a finding that correlates with the low expression of *gnb5* and consequently of Flr in this brain area.

The defects observed in the behavior of Flr mice could be explained by the deficient transport of synaptic proteins and endoplasmic reticulum (ER) to the PSD. Examining synaptosomes from Flr mice, we found that the Homer-IP3 receptor complex that binds to Shank and the MAGUKS (PSD95, PSD93 and SAP102) were significantly reduced (Figure 5). The IP3 receptor is integrated into the smooth endoplasmic reticulum (ER) membrane, enabling the MyoVa motor protein to pull the entire PSD-95 scaffolding complex with the smooth endoplasmic reticulum into spine synapses (Hammer and Sellers, 2011), thus explaining the deficient localization at the synapse.

Interestingly, SAP102, the earliest appearing MAGUK, was also reduced. SAP102 is not palmitoylated and therefore cannot travel by vesicular transport (El-Husseini et al., 2000). SAP102 is known to traffic to the synapse of young neurons in association with the NMDAR subunit GluN2B (Washbourne et al., 2004) but might not do so in older neurons such as the ones used in this study. GluN2B was not reduced in Flr relative to WT_FLR_ synaptosomes (Figure 5 D1-2). For this reason, we suggest that defects in synaptic protein synthesis might explain the decreased SAP102 presence at these older hippocampal synapses. SAP102 synthesis has been observed in isolated CA1 hippocampal neuropil (Cajigas et al., 2012). Defects in local protein synthesis also could account for some decreases in PSD-95, PSD-93 and other synaptic proteins we examined. We note that as techniques for isolating transcripts in neurons have improved more proteins have been found to be synthesized outside of the neuronal cell body in unexpected dendritic and axon compartments (Cajigas et al., 2012; Holt and Schuman, 2013).

Our previous work demonstrated that in mice homozygous for the Flr mutation visual cortical neurons have normal NMDAR-dependent LTP but lack NMDAR-dependent LTD (Yoshii et al., 2013). We also showed that in Flr mice many molecules critical to normal glutamate synaptic function are significantly reduced and misplaced throughout Flr dendritic shafts, including AMPA receptor (AMPAR) subunits, Dynamin-3 (critical for AMPAR) and other proteins that undergo endocytosis at synapses (Lu et al., 2007; Morimoto-Tomita et al., 2009; Petrini et al., 2009; Selvakumar et al., 2009; Kato et al., 2010). Our current study shows that in the hippocampus mGluR-dependent LTD was also diminished in Flr animals (Figure 6) and that positive modulators of mGluR did not revert this aspect of the Flr phenotype (Figure 7). These observations suggest that the reduced transport and localization of synaptic proteins in Flailer animals directly affects the induction of LTD, which cannot be recovered by the solely activation of mGluR. The lack of response to CDPPB was expected in Flr mice, because mGluR5-LTD requires protein synthesis that, in turn, requires long Homer in the glutamate scaffold (to maintain the protein synthesizing machinery near the receptor (Tu et al., 1999)) and as seen in Figure 5C Homer is significantly depleted in Flr synaptosomes.

Disruption of LTD has been widely associated with ASD in mutant mouse models of TSC1, Pcdh10, or Pten (Bateup et al., 2011; Tsai et al., 2012; Takeuchi et al., 2013). The lack of LTD in these mutant animals severely affect developmental activity-dependent synapse elimination or “pruning”. The absence of LTD and the consequent hyperconnectivity between neurons in transgenic mice carrying human ASD mutations, suggest that depression and elimination of early axon connections might be a highly vulnerable process in the developing brain. Understanding which connections are sensitive to disruption of this important event and how the disruption causes abnormal behavior will be important steps in mechanistically addressing psychiatric disease. We suggest that the Flailer mouse could significantly facilitate this endeavor. Indeed, much of the abnormal Flr behavior we describe as well as the differences in hippocampus synaptosome proteins between Flr vs WT_FLR_ mice can be related to the abnormally high level of synaptic function added by LTP but not removed by LTD in this mouse.

The abnormal behaviors of the Flr mutant mouse offer a unique opportunity to explore the importance of LTD and synaptic transport during normal development and behavior. Editing the Flr genome to inactivate at least one copy of the Flr gene should restore both normal MyoVA function and a WT phenotype (Bustos et al., 2015 SFN 390.23/C18). Then conditional knockouts specific to different neuron types could be used to identify the anatomical pathways and brain areas responsible for the abnormal behaviors of Flr mice. Such information might identify orthologous brain pathways disrupted in human disorders that present with abnormal behaviors comparable to those of Flr mice. Subsequently, studies of the *flr* gene could facilitate pharmaceutical or behavioral interventions to mitigate or cure some of the most devastating human neuropsychiatric diseases.

## EXPERIMENTAL PROCEDURES

### Animals

Flailer mice, line B6CBACa-A^*WJ*^/A, *tb*^*2J*^*/tb*^*2J*^ F_2_ from the frozen embryo repository at Jackson Laboratory (Bar Harbor, ME), were bred as described in Jones et al., (2000) and supplied to us as pairs from their BcC-flr congenic line by the laboratory of Professor Miriam Meisler, University of Michigan, Ann Arbor. Animals were rederived and maintained in the MIT IACUC. BcC-flr mice used for the current experiments were crossed with Thy-1 EGFP mice (Feng et al., 2000) to obtain F1 heterozygotes and subsequently bred to homozygosity for the *flr* gene. WT Flr-GFP animals were also obtained from these crosses. Both strains were bred for a subsequent 4 years in our laboratory. Genotypes were determined by PCR from mouse ear clippings (http://www.transnetyx.com/Services/Automated Genotyping.aspx). Animals were maintained at 23°C in the MIT ICACU facility with a 12 h light/dark cycle and food and water available *ad libitum*.

### Quantitative western blotting

Homogenate and synaptosome fractions were prepared from whole hippocampus tissue collected from P25-29 WT and Flr. Protein concentrations were determined with BCA assays (Pierce, IL). Proteins were electrophoretically separated on 4–12% SDS-PAGE gels and transferred to a PVDF membrane (Millipore, MA). Membranes were subsequently blocked with blocking buffer (Sigma, MO) and then incubated with primary antibodies overnight at 4°C, rinsed, and then incubated with secondary antibodies for 30 min at room temperature (RT). Blots were developed using chemiluminescence (Pierce. IL). Tubulin β III or β actin bands on the same blot were used as loading controls. Band intensities, were confirmed to be in the linear range and then quantified using Image J software. At least three separate protein samples were used for quantification.

### *In Vitro* Electrophysiology

Acute slices were prepared from either Flr or WT. Animals were anesthetized with isoflurane and decapitated. Brains were quickly removed and chilled in ice-cold dissection buffer. Transverse dorsal hippocampal 350-400 micron sections were cut using a VT-1000S vibratome (Leica, Germany) in ice cold carbogenated sucrose solution containing (in mM) 240 Sucrose, 2.5 KCl, 1 CaCl_2_, 5 MgSO_4_, 26 Na_2_HCO_3_, 1NaH_2_PO_3_, 11 Glucose, transferred to a chamber containing aerated artificial cerebrospinal fluid (ACSF) for 30 min at 32°C then maintained at RT (22-24°C) for a minimum of 1 hr prior to electrophysiology recordings. Electrodes were 4M NaCl (Sigma, MO) filled borosilicate glass (WPI Sarasota, FL) pulled to 1-3MΩ-tip resistance using a P-97 puller (Sutter Instruments, CA). For recordings, slices were submerged and perfused (3 ml/min) in a carbogen-saturated at RT. Stimulating electrodes were fabricated tungsten bipolar electrodes (WPI) driven by ISO-STIM-01D (NPI Electronic GmbH, Germany). Field potentials were recorded in stratum radiatum of CA1, amplified with a MultiClamp 700B Axopatch 200B, digitized with a Digidata 1440A, filtered at 2 kHz, sampled at 10 kHz and stored in a computer that provided both on--line display and off-line data analysis using pClamp 10.2 and Clampfit 10.2 software (Mol. Devices Corp., CA).

### Dendritic Protrusion density analysis

Male WT and Flr GFP mice were anesthetized with isoflurane and transcardially perfused with PBS followed by 4% paraformaldehyde (PFA) in PBS. Brains were dissected, post-fixed with 4% PFA at 4°C. The hippocampus was removed for transverse sectioning. Images were obtained by with a confocal microscope (Nikon PCM2000 MVI, Japan) using a 60x oil-immersion objective and zoom factor of 2 images. Multiple optical sections 0.5 µm were analysed using neurolucida Software (MBF Bioscience, VT).

#### Behavior Analysis

Adult male mice were used for all behavior analyses except for pup vocalizations. All observers were blind to the phenotype of the mouse unless otherwise stated.

#### Grooming behaviors

After 10 minutes of habituation to a fresh cage mice were video-taped for 24 h under 700 lux day lighting, (12 hr.) and ∼ 2 lux red light at night, (12 hr) illuminations. Grooming behaviors recorded at the onset of dark cycle (19:00–20:00 h) were studied using an automated home-cage behavioral phenotyping system (Jhuang et al., 2010) designed by the Poggio laboratory in the McGovern Institute with manual corrections. The videos shown in Supplementary S4A were taken in plexiglass cubicles to illustrate details in Flr grooming.

#### Elevated plus maze

We used a standard elevated plus maze as described in Komada et al., 2008. Each test session lasted 5 minutes. Scoring was accomplished using Ethovision XT Noldus Observer software.

### Three-Chamber social approach test

The three-chamber test for sociability was performed as previously described (Moy et al., 2004) with minor modifications. A WT mouse was placed in the central chamber separated by opaque panels with holes providing passage between chambers. A novel male mouse was held in a wired cage allowing visual, tactile, and olfactory contact. Video recordings of the introduced mouse were taken for the following 10 min period.

### Social Proximity Tests

Social proximity testing was conducted in a clear rectangular chamber constructed of acrylic plastic under dim red light as previously described (Defensor et al., 2011). For testing, two non-cagemate mice were simultaneously placed in the testing chamber for a 10 min trial. Either two mice of the same strain or one Flr and one WT were placed in the chamber. The entire test period was videotaped from two cameras providing front and side views. All behaviors involved some contact between the 2 animals.

### Morris Water maze

Before the spatial memory trials, mice were taught the location of a transparent lucite platform (10 × 10 cm) submerged just underneath the surface of the water. Memory for the platform location was assessed during four consecutive spatial trials 24hrs later. For all spatial trials, swim distance (m) and swim speed (m/s) were recorded. Flr and WT swim speed were not significantly different (data not shown).

### Fear Conditioning

Standard fear conditioning paradigms were used to test for freezing to context in standard fear-conditioning chambers housed in sound-attenuating boxes (MED Associates, St. Albans, VT). Twenty-four hours after a 5min placement in the chamber with a 180s noise (85db) stimulus followed immediately by a 2s (0.6mA) mild shock mice were placed back into the same chambers. Freezing to the context provided by the box alone was assessed over 5min. Cued stimulus fear conditioning was evaluated 2hrs later in altered chambers. Freezing in the absence of the cued stimulus was assessed during the first 180s of the test session, then the noise (85db) was presented and freezing to this cued stimulus was assessed.

### Communication Deficits

Pups (P2-P14) were isolated from their mother and littermates for 10 min. For testing, pups were gently placed into a glass isolation container containing clean bedding material and surrounded by a sound attenuating box (18×18×18 cm) made of 4mm thick Styrofoam. An ultrasound Gate system monitored USV emission. Acoustic data were recorded with a sampling rate of 250,000 Hz in 16-bit format by a recorder (version 2.97; Avisoft Bioacoustics, Germany). For acoustical analysis, recordings were transferred to Avisoft SASLab Pro (version 4.50; Avisoft Bioacoustics, Germany)

### Data Analysis

Statistical analyses were completed using paired Student’s t tests and Rank Sum test.

## Author Contributions

S.P. – Designed the work did all the behavior, imaging studies and drafted the initial copy of the manuscript.

Y.M – Did synaptosome western blot

JP. Z – Electrophysiological recordings

F.J.B. – revised the manuscript

C.T. – discovered the lack of mGluR5 –LTD in Flailer hippocampus

R.A. –S.P.’s Ph.D. advisor

M.C.P. – supervised and supported all aspects of the work and revised the manuscript.

## Acknowledgment

This work was supported by grants-R01-EY014074, R01-EY014420 to MCP. SP was supported by MIT-Portugal Program, the Portuguese Foundation for Science and Technology (SFRH/BD/33726/2009) as a graduate student and by the McGovern Institute for Brain Research as a postdoctoral fellow. FJB was a Simons center for the Social Brain Postdoctoral Fellow at MIT and a Pew Latin American Postdoctoral Fellow. FJB is funded by FONDECYT 11180540 and CONICYT-PAI 77180077. We thank Prof. Glen Prusky and Nazia Alam, Weill Cornell Medical College for helping us with the visual acuity test; Stanley Center, Prof. Mriganka Sur’s Lab and Prof. Gouping Feng’s Lab for behavioral equipment; Prof. Tomaso Poggio and Nicholas Edelman for development and setting up of the automated phenotyping system for mouse grooming behavior. We thank MCP lab members for reading the manuscript and helpful feedback.

